# Leveraging UMLS-driven NLP to enhance identification of influenza predictors derived from electronic medical record data

**DOI:** 10.1101/2020.04.24.058982

**Authors:** Kari A. Stephens, Margaret A. Au, Meliha Yetisgen, Barry Lutz, Monica Zigman Suchsland, Mark H. Ebell, Matthew Thompson

## Abstract

**Objective:** Multiple clinical prediction rules have been developed, but lack validation. This study aims to identify a set of prediction algorithms for influenza, based on electronic health record (EHR) structured data and clinical notes derived data using Unified Medical Language System (UMLS) driven natural language processing (NLP).

**Materials and Methods:** Data were extracted from an enterprise-wide data warehouse for all patients who tested positive for influenza and were seen in ambulatory care between 2009 and 2019 (*N* = 7,278). A text processing pipeline was used to analyze chart notes for UMLS terms for symptoms of interest to improve data quality completeness. Three models, which step up complexity of the dataset and predictors, were tested with least absolute shrinkage and selection operator (LASSO)-selected parameters to identify predictors for influenza. Receiver operating characteristic (ROC) curves compared test accuracy across the three models.

**Results:** Three models identified 7, 8, and 10 predictors, and the most complex model performed best. The addition of the UMLS-driven NLP symptoms data improved data quality (false negatives) and increased the number of significant predictors. NLP also increased the strength of the models, as did the addition of two-way predictor interactions.

**Discussion:** The EHR is a feasible source for offering rapidly accessible datasets for influenza related prediction research that was used to produce a prediction model for influenza. Combining data collected in routine care with data science methods improved a prediction model for influenza, and in the future, could be used to drive diagnostics at the point of care.

## INTRODUCTION

Multiple clinical prediction rules (CPRs) have been developed that aim to improve the evidence supporting diagnostic and predictive decision in clinical settings.[1] CPRs typically aim to provide an evidence-based tool that can be used to improve clinical diagnostic or predictive decision making processes, and many have been widely adopted into routine care (e.g., related to cardiovascular disease, respiratory, and musculoskeletal conditions).[2] As the numbers of CPRs being developed has increased, guidelines and standards have been developed that outline the steps needed to support implementation into routine clinical practice[3] such as external validation, assessing impact on clinical decision making and patient outcomes.[4]

Most CPRs are derived and validated using data that has been prospectively collected directly from patients in the context of a research study. These data typically involve time and labor intensive standardized collection of key predictor variables, such as clinical signs and symptoms, as well as standardized methods to ascertain outcomes. Obtaining suitable data direct from patient populations in multiple settings involves considerable research infrastructure and support, limiting rapid validation and implementation. Relatively few CPRs have been evaluated beyond initial pilot development, due to insufficient data and difficulty with access.[2, 4] In fact, in primary care, a review of 434 CPRs showed that only 55% had been validated.[2]

Data routinely collected in care, through the electronic health record (EHR), can potentially be used to validate CPRs, reducing the need to conduct additional costly prospective studies. With the widespread use of EHR systems to record and store clinical data collected in routine care, large datasets in enterprise wide data repositories may provide a powerful source of validation data for emerging CPRs. EHR data are widely available and have rich breadth across symptoms, diagnoses, and other clinical findings.

A major known EHR data quality limitation includes variable non-random rates of missing values,[5] particularly among structured or coded data elements in the EHR. Natural language processing (NLP) methods provide the means to rapidly interrogate and derive structured data out of clinical notes to improve non-random missing data values.[6] The Unified Medical Language System (UMLS) Metathesaurus provides a rich set of medical terminology that can be used in combination with NLP methods to derive structured clinical concepts rapidly from notes. Yetisgen and colleagues developed a text processing pipeline for extracting all known UMLS concepts with surrounding semantics from any clinical note captured in an institutional enterprise wide data warehouse.[7] These UMLS NLP derived structured data can enable capture, for example, of provider identified symptoms charted as clinical text, rather than as an International Classification of Diseases (ICD) code (e.g., cough, fever).

Detection of infectious diseases offer a prime area to explore utility of CPRs that can be derived and validated rapidly from EHR data and UMLS enabled NLP methods because of their pervasiveness in the population and the potential benefits of CPRs for rapid diagnosis. For example, multiple influenza related CPRs mainly derived from resource intensive patient reported symptoms have been developed, but lack validation.[8, 9] Influenza CPRs provide a valuable use case for testing the utility of EHR data for development and validation of CPRs. Patients with influenza who present for care are often tested using a polymerase chain reaction (PCR) lab test, offering an accessible EHR derived gold standard outcome, and have symptoms documented by providers using ICD codes and text within clinical notes. In addition, patients with influenza commonly present for care with influenza-like illness related symptoms that have been well defined based on previous observational studies.[10]

This study aims to evaluate a set of prediction algorithms for influenza, based on EHR structured data and clinical notes derived data using a UMLS-driven NLP method, to determine the utility of EHR data for providing a rapidly re-usable large dataset for developing and validating CPRs. These EHR prediction algorithms are hypothesized to improve CPR breadth and strength through the addition of NLP derived data and use of machine learning methods to explore interactions between predictors. UMLS-driven NLP methods are expected to improve rates of false negatives in symptom data detection, due to lack of coded data (missing data from provider entry), and improve CPRs that do not use data from clinical notes.

## METHODS

### EHR Dataset

De-identified data were obtained from University of Washington Medicine’s enterprise wide data repository (EDW), which stores EHR data for over four million patients, with IRB approval granted via Human Subjects Assurance FWA #00006878, IRB ID STUDY00008069. PCR testing for influenza was recorded on 22,938 patients between 2009 and 2019, of whom 13,898 patients had this test ordered from an ambulatory care clinic setting. Patients who were missing at least one measure of vitals data, specifically either heart rate, blood pressure (systolic and diastolic), or temperature, were excluded. Sensitivity analyses between patients with and without complete vitals data showed no significant differences. All patients with a PCR test, seen in an ambulatory care clinic setting, and with complete vitals data (*N* = 7,278) were used in the final analyses.

#### Influenza “Gold Standard” Outcome

A positive influenza PCR test was used as ground truth for a predicted outcome of a “true” test of influenza, and were identified via lab data within the EDW (see attached appendix for proprietary lab codes). A set of 27 lab tests was defined through an iterative process that began with identifying each test with the text “influenza” in the description. That list was then validated by a group of context experts on the research team. PCR tests were classified as “positive” based on the raw result from the lab medicine system indicating it as “positive.”

#### Influenza Predictors

Influenza predictors were identified through predictors noted in published CPRs [9] and through domain expert input by both infectious disease and primary care providers. The following predictors were identified as possible structured (i.e., coded fields) and unstructured (i.e., clinical notes with embedded UMLS concepts) data to be extracted from the EHR. Patient demographics and social history predictors included age, gender, race, ethnicity, public insurance status (yes/no), and smoking status (known at the time of the patient’s PCR test). The PCR test date was used to identify ambulatory care visits with documented patient symptoms and vitals from data captured in the EHR. Influenza vaccination status was identified if given within six months of the PCR test date. Patient symptom predictors were all designated dichotomously as present versus not present across 16 symptoms that included fever, sort throat, cough, myalgia, crepitations, dyspnea/shortness of breath, coryza/nasal decongestion, hemoptysis, myalgia, chills/rigors/sweating, malaise/fatigue/weakness, headache, diarrhea, vomiting, lack of appetite, and rash and ear pain/discharge. Symptoms were identified using both ICD-9 and −10 codes and UMLS concepts extracted using NLP (see attached appendix). Symptoms were classified as dichotomously present or not present at the time the PCR test was ordered. Patient vitals data included numeric values of heart rate, blood pressure, and temperature – which was an additional predictor to fever, noted above as a dichotomous symptom to allow for more specificity of the symptom via a numeric value.

#### UMLS-driven Natural Language Processing (NLP) Derived Influenza Symptom Predictors

This study adapted a text processing pipeline, which used NLP to analyze chart notes from the index visit for UMLS terms for symptoms of interest.[7] The NLP pipeline is used to process all clinical notes stored in EDW. The pipeline first chunked each clinical note into sentences with OpenNLP sentence chunker,[11] next it extracted mentions of UMLS Metathesaurus concepts with their associated assertion values. UMLS concept extraction was done with a tool developed by National Library of Medicine (NLM) called MetaMap.[12] A lightweight Java implementation (Metamap Lite) was used in our pipeline due to processing speed and ease of use. In a recent study, MetaMap Lite demonstrated real-time speed and extraction performance comparable to or exceeding those of MetaMap and other popular biomedical text processing tools,[13] clinical Text Analysis and Knowledge Extraction System (cTAKES),[14] and DNorm.[15] Metamap-Lite extracted medical problems, tests, and treatments from 2010 i2b2 concepts dataset with precision 47.0, recall 31.9, and F1 38.0.[13]

After identifying the UMLS concepts, the NLP pipeline assigned each extracted UMLS Metathesaurus concept an assertion value (*present, absent, conditional, hypothetical, possible, not-patient*) with an in-house statistical assertion classifier. While building the in-house assertion classifer, the Stanford NLP library[16] was used for tokenization, POS tagging, and dependency parsing to capture a wide range of syntactic and semantic features presented in clinical text. Those features were used to train the SVM based state-of-the-art assertion classifier. Our state-of-the-art assertion classifier produced Micro-F1 94.23 when trained and tested on the i2b2 2010 assertion dataset detailed elsewhere.[17]

The extracted information of UMLS concepts with assigned assertion values as well as character indexes of the identified concepts within notes are stored in a database table to be used as a semantic index for clinical notes within the EDW. In this study, the semantic index was queried to identify patients with mentions of influenza symptoms. We first identified the UMLS concepts for each influenza symptom and searched the semantic index for those concepts with assertion value *present*. We evaluated the performance of the NLP pipeline for symptoms that were found to be present in more than 5% of the overall cohort (e.g., terms included chills, congestion, cough, diarrhea, fatigue, fever, myalgia, pharyngitis, shortness of breath, sore throat and vomiting). Twenty randomly selected notes were extracted and used to check the correctness of a total of 200 symptom extractions with their associated assertion values. Our analysis indicated the NLP pipeline successfully identified the symptoms with assertion values with 0.89 precision. All 17 negation cases (e.g., denies fever) were identified correctly. Assertion value for the symptom for 14 cases could not be determined. The present assertion value was confused across cases with hypothetical (5 cases), conditional (1 case), and not patient (1 case).

### Analyses

Comparisons between patient characteristics in those testing positive versus negative for influenza were assessed using chi square or Fisher’s Exact tests. The patient cohort was split into a 70% training dataset and 30% validation dataset, in order to develop and validate predictive models for PCR-detected influenza. Variable selection and regularization, using a least absolute shrinkage and selection operator (LASSO) regression approach, was used to test prediction models, and Schwarz’ Bayesian Criterion (SBC) was used to compare model performance. Minimization of SBC was the criteria used to select final models and to determine optimal penalty coefficient to be 0.01. [18] Three LASSO predictive models were constructed: 1) Model 1 – all predictors as single prediction factors with (ICD) coded symptoms, 2) Model 2 – same as Model 1 with ICD symptoms data augmented by UMLS-driven NLP derived data, and 3) Model 3 – same as Model 2 combined with all two-way categorical interactions added as additional predictors. Logistic regression models with the LASSO-selected parameters (and the associated main effects in the case of interactions) were calculated on the validation data to summarize selected candidate variables’ associations with the outcome. Receiver operating characteristic (ROC) curves were used to illustrate sensitivity and specificity of the predictions for the three models, to compare the benefit of each model. Receiver operating characteristic (ROC) curves were used to illustrate sensitivity and specificity of the predictions for the three models.

## RESULTS

### Patient Characteristics and Symptoms

Patients with a positive versus negative PCR influenza test differed significantly on all patient characteristics except gender (see Table 1). Patients with a positive PCR test had a lower proportion with public insurance, were slightly less likely to be a smoker, more likely to be younger and to have been vaccinated for influenza, and less likely to be White than those with a negative test.

**Table 1.**
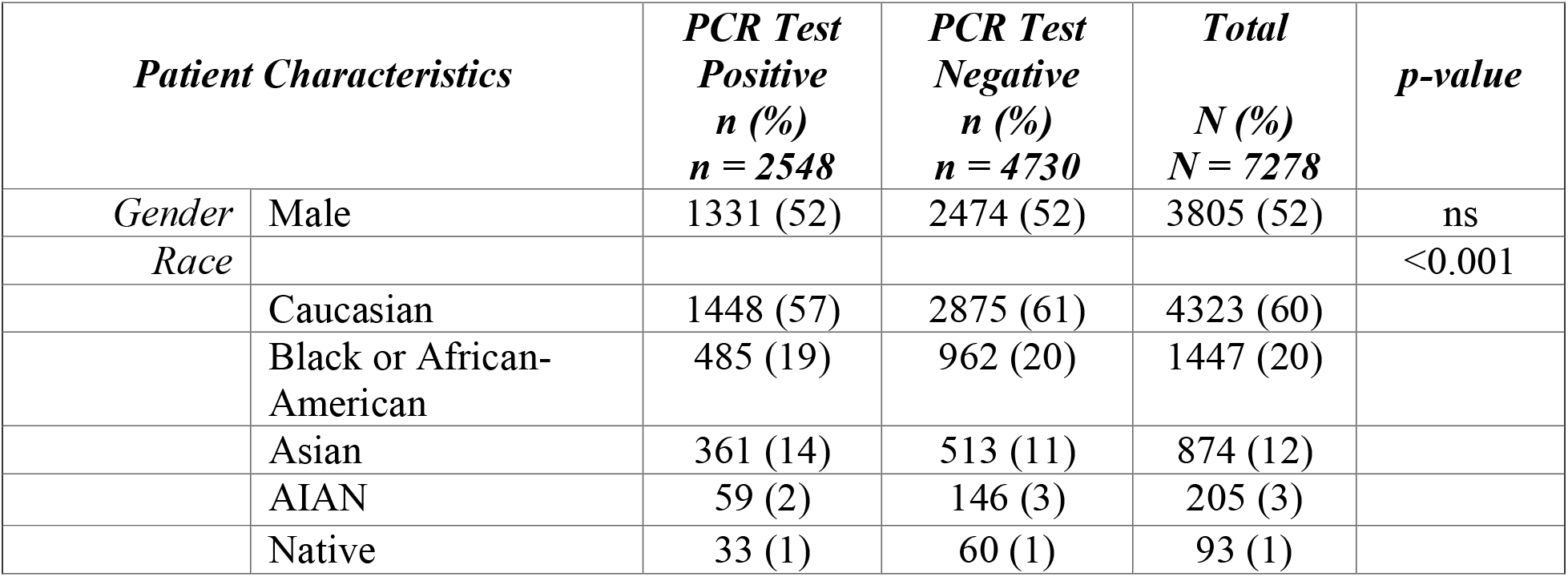
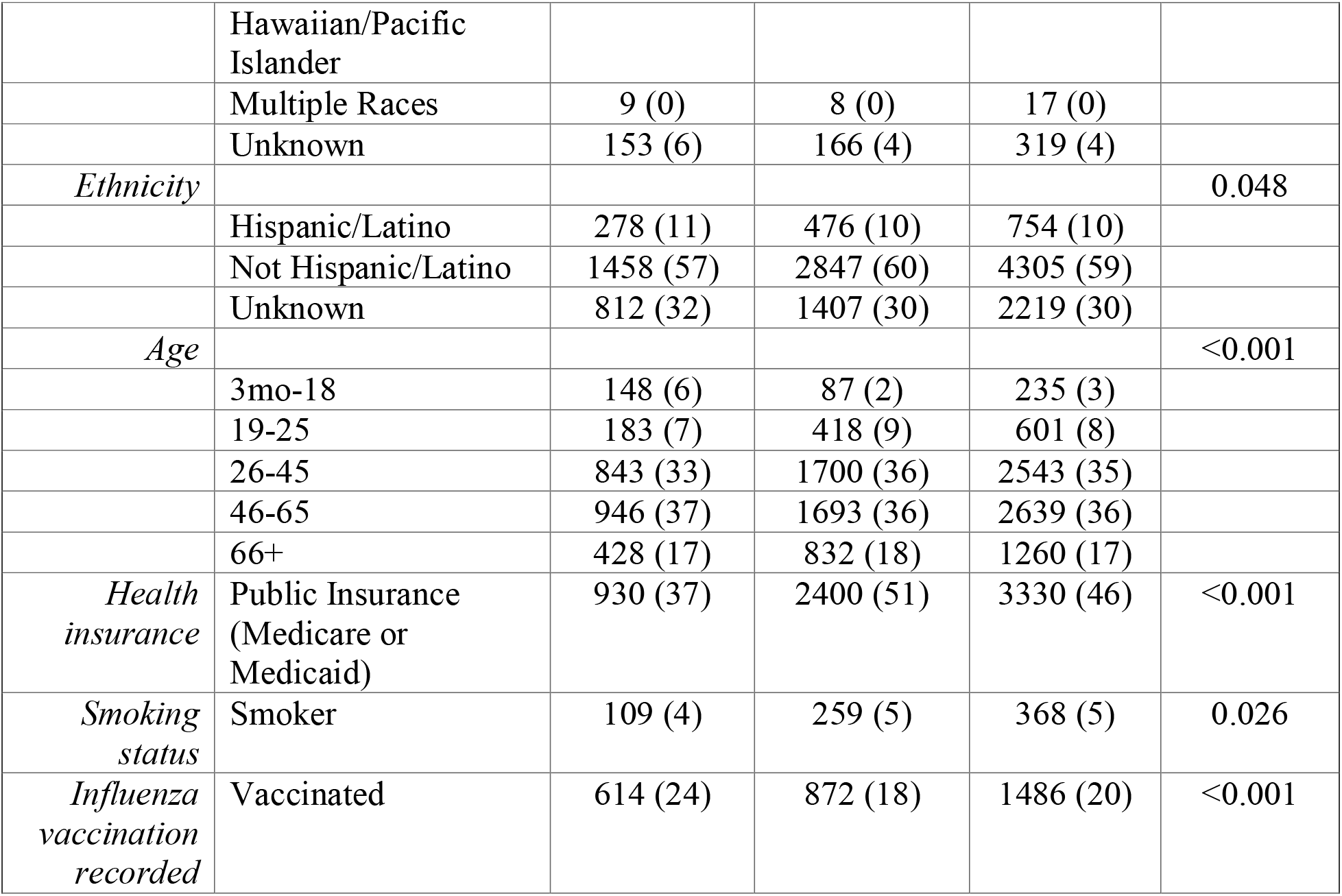
Patient characteristics for patients that had a PCR test in an outpatient clinic setting between 2009 and 2019.

Patients with a positive PCR test were more likely to have symptoms that have previously been associated with influenza (e.g., fever, sore throat, cough, myalgia, chills/sweats, nasal congestion) than patients with a negative PCR test (see Table 2). Influenza-related symptom detection rates among patients with PCR tests varied greatly, with some symptoms (e.g., fever, cough/expectoration) being found in over half of patients (see Table 2). Extraction of symptoms via UMLS-driven NLP derived data increased detection across 10 symptoms (i.e., 8-23% for patients with a positive PCR test, 8-18% for patients with a negative PCR test).

**Table 2.**
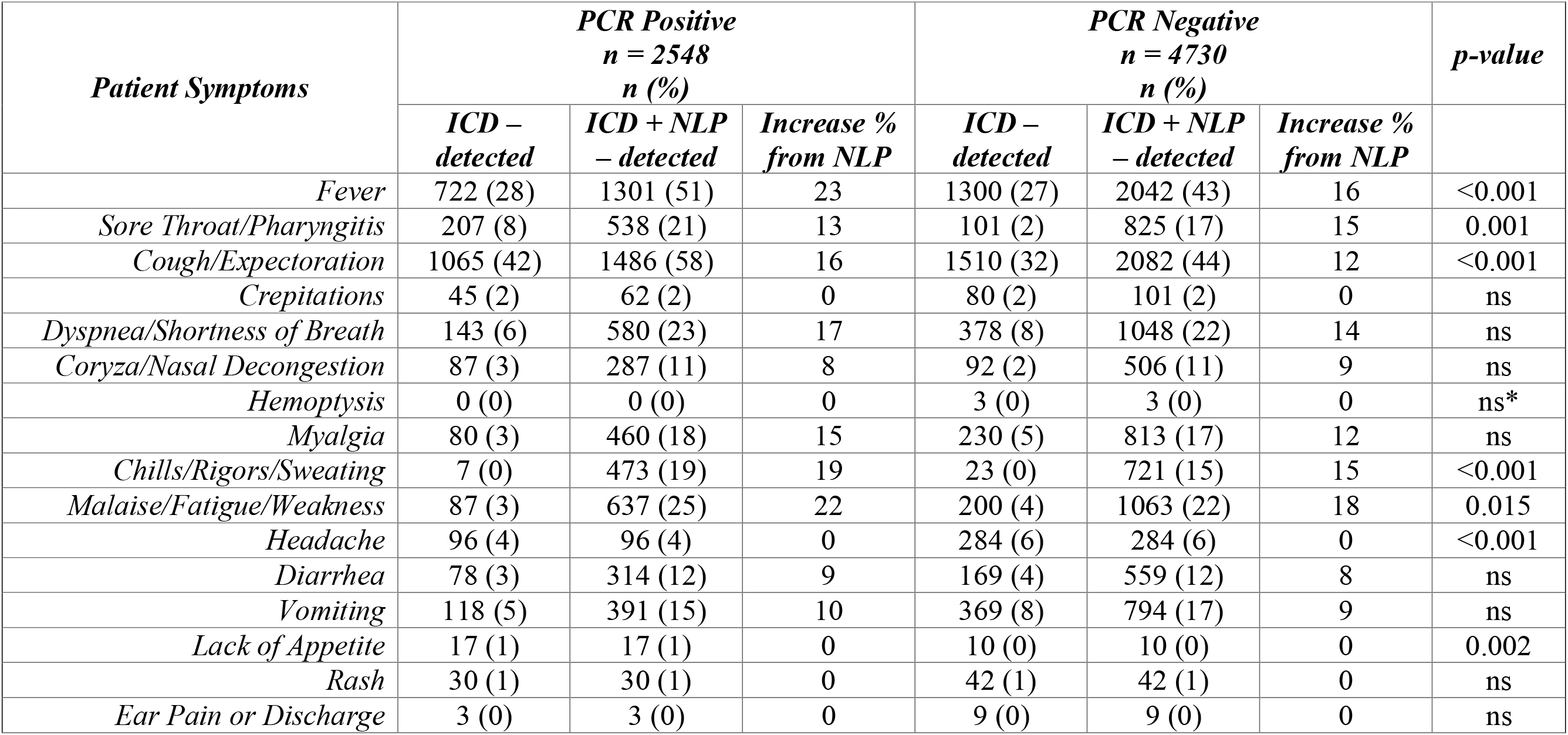
Influenza related symptom detection differences for patients with a positive vs. negative PCR test.

### LASSO Regression Models

The models identified increasingly higher numbers of predictors as they added in UMLS-derived NLP data and evaluated two-way interactions (see Tables 3 and 4). Model 1 selected seven predictors using LASSO regression in the training dataset. Model 2, with the addition of UMLS-derived NLP symptom data, selected the same set of predictors as Model 1 and added a new symptom predictor of fever. Logistic regression performed on the validation data demonstrated a modest, but significant, 2% improvement in AUC from Model 1 (0.66) to Model 2 (0.68), *p* < 0.001 (see Figure 1). Model 3 (AUC = 0.71), with the addition of all two-way interactions between categorical predictors, also modestly, but significantly increased from Models 1 and 2 (*p*’s < 0.001). Model 3 selected the same demographics and vitals predictors as Model 2 and added four new predictor interactions (i.e., public insurance*cough, fever*cough, current vaccination*cough, and public insurance*myalgia). The logistic regression model including all the selected effects, along with any main effects for the selected interactions, showed a 3% additional significant improvement in the AUC from Model 2 and 5% from Model 1. While cough, public insurance, current vaccination, and temperature were significant predictors in the validation data for all three models, the presence of cough was associated with decreased odds of influenza and current vaccination was associated with increased odds of influenza (see Table 4). In Model 3, the interactions of public insurance*cough, fever*cough, and public insurance*myalgia were also highly significant in the validation data. The presence of significant interactions between individual factors in the model suggests that the associations of individual factors with influenza vary across values of other patient factors.

**Table 3.** Predictors compared across LASSO models derived from EHR data vs. a clinical prediction rule, Flu Score, derived from direct patient data.

| *Patient Predictors* | *Model 1:*<br>*LASSO*<br>*with ICD* | *Model 2:*<br>*LASSO*<br>*with ICD*<br>*+ UMLS NLP* | *Model 3:*<br>*LASSO*<br>*with ICD*<br>*+ UMLS NLP*<br>*&*<br>*interactions* | *Flu*<br>*Score 3*<br>*CPR[9]* |
| --- | --- | --- | --- | --- |
| **Characteristics** |  |  |  |  |
| Gender | ns | ns | ns | -- |
| Race | ns | ns | ns | -- |
| Ethnicity | ns | ns | ns | -- |
| Age | ns | ns | ns | -- |
| Public Health Insurance | *** | *** | *** | -- |
| Smoking Status | ns | ns | ns | -- |
| Current Vaccination | *** | *** | *** | ns |
| **Symptoms** |  |  |  |  |
| Fever | ns | *** | ns | ***<sup>††</sup> |
| Sore Throat/Pharyngitis | ns | ns | ns | ns |
| Cough/Expectoration | *** | *** | ns | ***<sup>††</sup> |
| Crepitations | ns | ns | ns | -- |
| Dyspnea/Shortness of Breath | ns | ns | ns | -- |
| Coryza/Nasal Congestion | ns | ns | ns | -- |
| Hemoptysis | ns | ns | ns | -- |
| Myalgia | ns | ns | ns | *** |
| Chills/Rigors/Sweating | ns | ns | ns | *** |
| Malaise/Fatigue/Weakness | ns | ns | ns | -- |
| Headache | ns | ns | ns | -- |
| Diarrhea | ns | ns | ns | -- |
| Vomiting | ns | ns | ns | -- |
| Lack of Appetite | ns | ns | ns | -- |
| Rash | ns | ns | ns | -- |
| Ear Pain or Discharge | ns | ns | ns | -- |
| Onset < 48 Hours | -- | -- | -- | *** |
| **Vitals** |  |  |  |  |
| Heart Rate | *** | *** | *** | -- |
| Systolic Blood Pressure | *** | *** | *** | -- |
| Diastolic Blood Pressure | *** | *** | *** | -- |
| Temperature | *** | *** | *** | -- |
| **Interactions<sup>†</sup>** |  |  |  |  |
| Public Insurance*Cough | -- | -- | *** | -- |
| Fever*Cough | -- | -- | *** | -- |
| Current Vaccination*Cough | -- | -- | *** | -- |
| Public Insurance*Myalgia | -- | -- | *** | -- |

Note: \*\*\* = significant predictor; ns = not a significant predictor; -- = not available in the dataset for the model; <sup>†</sup> = non-significant two-way interactions not listed; <sup>††</sup> = fever and cough were combined as one predictor in Flu Score; in sensitivity analyses, the same categorical predictors are retained when vitals are removed from this cohort and from a cohort with incomplete vitals

**Table 4.**
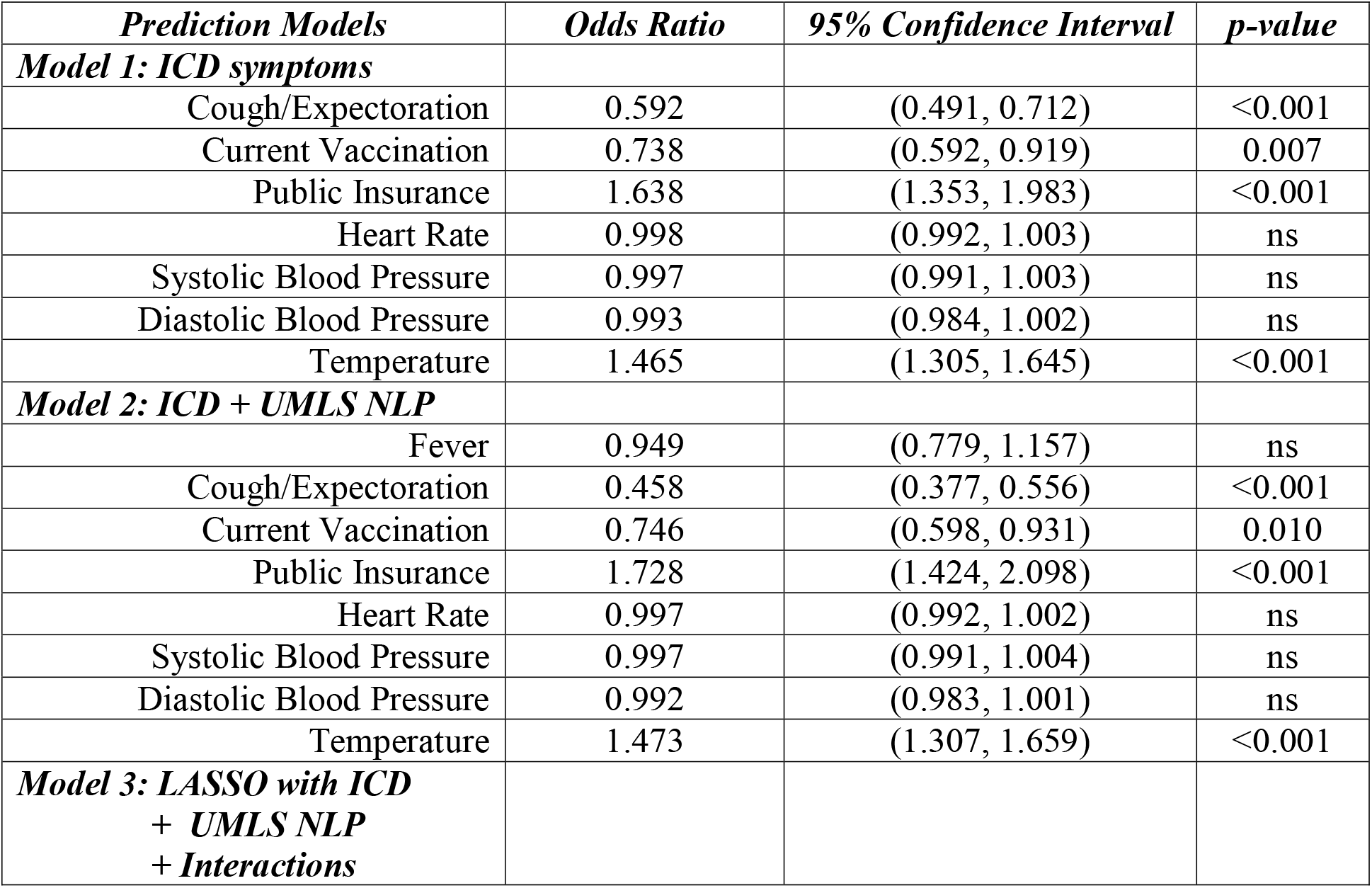
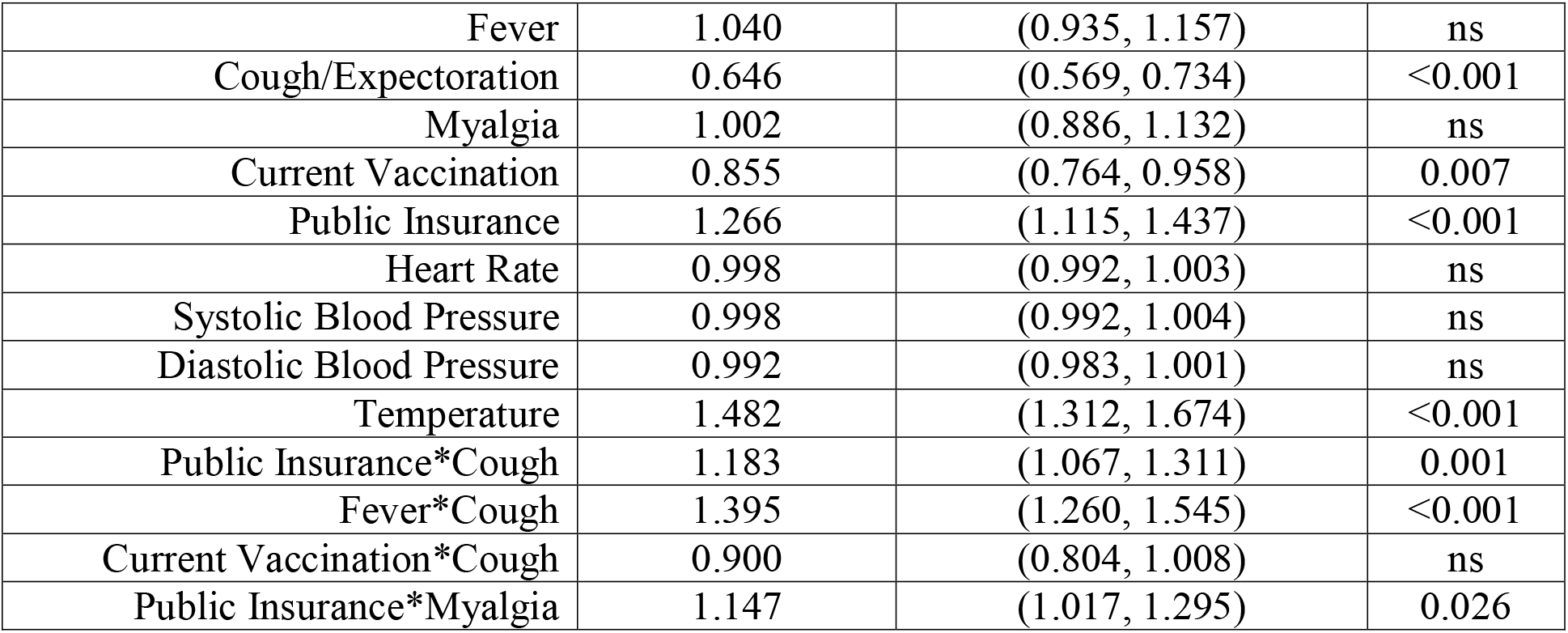
Results of logistic regression for influenza models tested from EHR derived data.

| **Prediction Models** | **Odds Ratio** | **95% Confidence Interval** | **p-value** |
| --- | --- | --- | --- |
| **Model 1: ICD symptoms** |  |  |  |
| Cough/Expectoration | 0.592 | (0.491, 0.712) | <0.001 |
| Current Vaccination | 0.738 | (0.592, 0.919) | 0.007 |
| Public Insurance | 1.638 | (1.353, 1.983) | <0.001 |
| Heart Rate | 0.998 | (0.992, 1.003) | ns |
| Systolic Blood Pressure | 0.997 | (0.991, 1.003) | ns |
| Diastolic Blood Pressure | 0.993 | (0.984, 1.002) | ns |
| Temperature | 1.465 | (1.305, 1.645) | <0.001 |
| **Model 2: ICD + UMLS NLP** |  |  |  |
| Fever | 0.949 | (0.779, 1.157) | ns |
| Cough/Expectoration | 0.458 | (0.377, 0.556) | <0.001 |
| Current Vaccination | 0.746 | (0.598, 0.931) | 0.010 |
| Public Insurance | 1.728 | (1.424, 2.098) | <0.001 |
| Heart Rate | 0.997 | (0.992, 1.002) | ns |
| Systolic Blood Pressure | 0.997 | (0.991, 1.004) | ns |
| Diastolic Blood Pressure | 0.992 | (0.983, 1.001) | ns |
| Temperature | 1.473 | (1.307, 1.659) | <0.001 |
| **Model 3: LASSO with ICD**<br>**+ UMLS NLP**<br>**+ Interactions** |  |  |  |
| Fever | 1.040 | (0.935, 1.157) | ns |
| Cough/Expectoration | 0.646 | (0.569, 0.734) | <0.001 |
| Myalgia | 1.002 | (0.886, 1.132) | ns |
| Current Vaccination | 0.855 | (0.764, 0.958) | 0.007 |
| Public Insurance | 1.266 | (1.115, 1.437) | <0.001 |
| Heart Rate | 0.998 | (0.992, 1.003) | ns |
| Systolic Blood Pressure | 0.998 | (0.992, 1.004) | ns |
| Diastolic Blood Pressure | 0.992 | (0.983, 1.001) | ns |
| Temperature | 1.482 | (1.312, 1.674) | <0.001 |
| Public Insurance*Cough | 1.183 | (1.067, 1.311) | 0.001 |
| Fever*Cough | 1.395 | (1.260, 1.545) | <0.001 |
| Current Vaccination*Cough | 0.900 | (0.804, 1.008) | ns |
| Public Insurance*Myalgia | 1.147 | (1.017, 1.295) | 0.026 |

**Figure 1.**
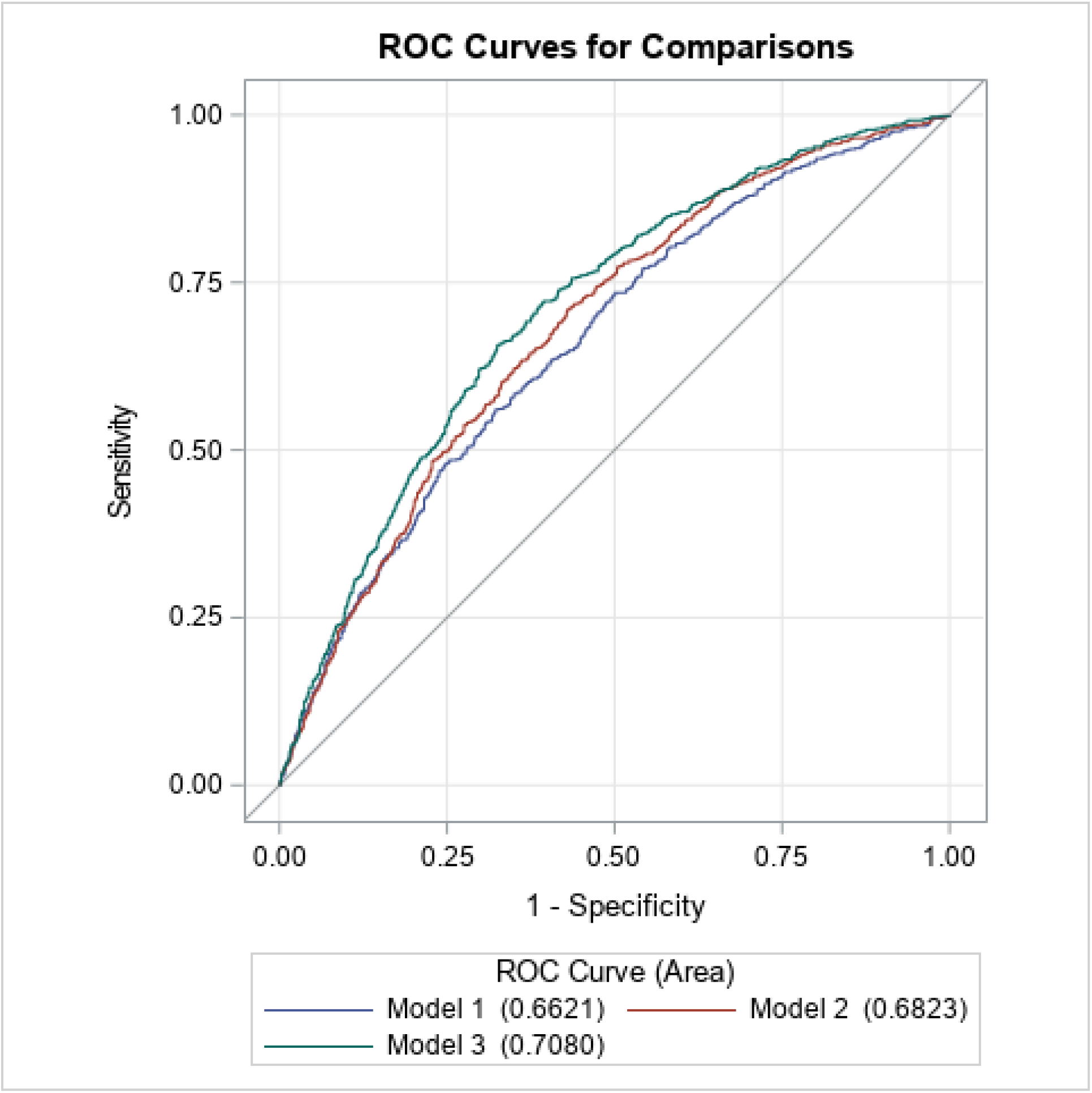
ROC curves across LASSO EHR derived models.

The Flu Score CPR examined data collected directly from patients and included six predictors as single factors in their model.[9] The models in this study examined data extracted from the EHR and included 30 predictors, which overlapped with five out of six predictors from the Flue Score CPR (see Table 3). The EHR derived data was not able provide onset time of symptoms (i.e., < 48 hours) to replicate Flu Score’s full set of predictors, which was notably significant in the Flu Score CPR model.

## DISCUSSION

EHR data were used to effectively create a prediction model by identifying a set of clear predictors that improved with the addition of UMLS-derived NLP symptom data by identifying more unique predictors and improving the overall performance of algorithms for predicting influenza in patients presenting in ambulatory care clinic settings. The EHR data offered several advances in predictive modeling of influenza including rapid access to a large cohort of patients, an expanded set of predictor data, and a rapid method to derive and validate several prediction algorithms.

The ease of extracting data from the EHR through pre-existing enterprise-wide warehouses within healthcare systems can fast track the derivation and validation of CPRs when cohorts can be easily identified, and predictors and outcome phenotypes can be clearly defined computationally. Patients at risk for influenza who presented with flu-like symptoms and who were potentially at risk of influenza infection were easily identified in the EHR based on the record of an order of a PCR test for influenza, and the positive test result allowed for a clear and objective reference standard outcome.

The EHR data allowed for examination of an expanded set of predictors that included demographics, vaccination, and vital signs collected routinely in care that are less feasible to collect directly from patient self-report via traditional methods like surveys, and are highly labor-intensive to collect prospectively over multiple clinic settings and multiple years. However, some kinds of predictors (i.e., vital signs of heart rate, activity levels) may now be detected by wearables and hold promise for potentially expanding the use of such predictors as part of early mHealth driven disease detection systems.

Influenza vaccination was associated with higher risk of infection, which seems counter intuitive, but the cohort of patients with a positive PCR test received significantly more vaccinations, this may be more of an indicator of service provision. Conducting research to identify new CPRs must account for patterns in datasets that might cause biases in samples based on service utilization patterns.

EHR datasets can be enriched by using NLP that leverages UMLS concepts to derive structured data out of unstructured clinical notes, which can improve data quality by reducing non-random missing data. In this study, the use of UMLS with NLP improved data quality, through the reduction of false negatives, and resulted in a modest increase in model performance. The text processing pipeline leveraged in this study can provide a reproducible method for deriving many UMLS concepts from clinical notes rapidly and with adequate precision. This provides a method to rapidly and feasibly create analytic-ready datasets without the time and resource-intensive prospective clinical data collection protocols often employed in development and validation of CPRs.

Potential limitations to this study include a lack of generalizability, given data were included from only one healthcare system, ambulatory care settings, and from only patients who received formal testing for influenza. However, this study could be replicated using data from other clinical settings or from individuals at home using mobile devices. This study also provides evidence that routine data collected in care, leveraging NLP methods, can be used to create clinical prediction algorithms, similar to public health surveillance related studies which have focused on overlapping subsets of variables we used.[19, 20] Data completeness may also have been an issue, given detection of symptom data, namely for fever, sore throat, cough, and myalgia, was lower than that previously found in the U.S. and Switzerland.[9] This may be a reflection of the sample bias or reflect a higher rate of false negatives (i.e., symptoms not recorded in the EHR despite being present) in the EHR data. While use of NLP strategies to reduce false negatives is important, rates of provider documentation should be evaluated.

Future directions can include embedding these prediction algorithms into clinical care, with data science innovations that can use routine data to drive prediction likelihood in real time for infectious disease diagnosis. In the immediate term, further validation of this algorithm is needed, using multi-site datasets across institutions to not only validate these findings, but also explore subgroup differences in prediction based on the large numbers that can be pooled from data collected in routine care and determine the nature of the relationship of variation in vitals (i.e., temperature, blood pressure, and heart rate) values to influenza. Deep learning algorithms, beyond LASSO methods, could be used to detect more powerful prediction as well. EHRs may provide a rich and easily accessible data source for conducting prediction data science driven research to study pandemics and prevent spread of disease.

The EHR is a feasible source for offering rapidly accessible datasets for influenza related prediction research that was used to produce a prediction model for influenza. Combining a rich set of data collected in routine care with data science methods that used NLP in combination with UMLS concepts improved data quality and the performance of machine learning prediction methods. Data collected in routine care can be combined with data science methods to help speed innovations in multiple areas of prediction and diagnosis.

## Supporting information

ICD and CUI codes

PCR lab codes

## Contributorship Statement

The authors contributed to the manuscript as follows – drafting lead KAS; design of the work KAS, MJT, BL, MAA; analysis MAA, KAS, MY; interpretation KAS, MAA, MT, BL, MHE, MZS; revisions, final approval ALL AUTHORS; accountable for all aspects of the work KAS, MT, BL

