## Supplementary material for "Leveraging UMLS-driven NLP to enhance identification of influenza predictors derived from electronic medical record data": ICD and CUI codes

| Symptom | ICD-9 Codes | ICD-10 Codes | UMLS Concept Codes |
|---|---|---|---|
| dyspnea (shortness of breath) | 786.09<br>786.05 | R06.00<br>R06.02 | C0013404 |
| expectoration (cough) | 786.2 | R05 | C0010200 |
| fever | 780.6 | R50.9 | C0015967<br>C0424755 |
| malaise / fatigue / weakness | 780.79 | R53.1<br>R53.8<br>R53.81<br>R53.83 | C0015672 |
| crepitations | 786.07<br>786.7<br>786.9 | R06.2<br>R09.89 | C0034642 |
| chills / rigors / sweating | 780.64 | R68.83 | C0085593<br>C0424790<br>C0038990 |
| myalgia | 729.1 | M79.1<br>R52 | C0231528 |
| hemoptysis | 786.3 | R04.2 | C0019079 |
| confined to bed | V49.84 | Z74.01 | not applicable |
| headache | 784 | R51 | C0018681 |
| pharyngitis / sore throat | 784.1<br>462 | J02<br>J02.9<br>R07.0 | C0031350<br>C0242429 |
| coryza / nasal congestion | 460<br>478.19 | R09.81<br>R09.82<br>J00<br>J34.89 | C0700148 |
| diarrhea | 787.91 | R19.7 | C0011991 |
| vomiting | 536.2<br>787.03<br>787.01 | R11.1<br>R11.10<br>R11.11<br>R11.12<br>R11.2 | C0042963 |
| nausea | 787.02<br>787.01 | R11.0<br>R11.2 | C0027497 |
| lack of appetite | 783 | R63.0 | C0003123 |
