## Supplementary material for "Leveraging UMLS-driven NLP to enhance identification of influenza predictors derived from electronic medical record data": PCR lab codes

| TestName |
| --- |
| Rapid Influenza B PCR Result |
| Seasonal H1N1 influenza A |
| Rapid Influenza A PCR Result |
| INFLUENZA SUBTYPING PANEL PCR |
| Influenza B PCR Screen |
| EPIDEMIC INFLUENZA PCR PANEL |
| Influenza A PCR Screen |
| INFLUENZA A; B AND RSV; RAPIC PCR |
| INFLUENZA A; B AND RSV; RAPID PCR |
| INFLUENZA A/B PCR (NWH) |
| INFLUENZA A/B & RSV BY PCR |
| Seasonal H3N2 influenza A |
| RAPID INFLUENZA A; B; RSV PCR |
| Influenza A Ct Value |
| RAPID INFLUENZA A; B; RSV PCR (NWH ONLY) |
| 2009 swine H1N1 influenza A |
| Influenza A PCR Qual Result |
| Influenza A; Rapid |
| INFLUENZA A/B AND RSV PCR WITH SUBTYPING |
| INFLUENZA A;B;RSV_RAPID PCR |
| RAPID INFLUENZA A AND B PCR |
| Influenza B PCR Qual Result |
| Influenza B Ct Value |
| Influenza B; Rapid |
| Influenza A/B PCR Result |
| INFLUENZA A/B AND RSV PANEL |
| ROUTINE INFLUENZA A/B & RSV |
